# The Role of the Subthalamic Nucleus in Inhibitory Control of Oculomotor Behavior in Parkinson’s Disease

**DOI:** 10.1101/606897

**Authors:** Shahab Bakhtiari, Ayca Altinkaya, Christopher C. Pack, Abbas F. Sadikot

## Abstract

The ability to inhibit an inappropriate action in a context is an important part of the human cognitive repertoire, and deficiencies in this ability are common in neurological and psychiatric disorders. An anti-saccade is a simple experimental task within the oculomotor repertoire that can be used to test this ability. The task involves an inhibition of a saccade to the peripheral target (pro-saccade) and generation of a voluntary eye movement toward the mirror position (anti-saccade). Previous studies provide evidence for a possible contribution from the basal ganglia in anti-saccade behavior. However, the precise role of different components in generation of anti-saccade behavior is still uncertain. Parkinson’s disease patients with implanted deep brain stimulation (DBS) in subthalamic nucleus (STN) provide us with a unique opportunity to investigate the role of STN in anti-saccade behavior. Previous attempts to show the effect of STN DBS on anti-saccades have produced conflicting observations. For example, the effect of STN DBS on anti-saccade error rate is not yet clear. Part of this inconsistency may be related to differences in dopaminergic states in different studies. Here, we tested Parkinson’s disease patients on anti- and pro-saccade tasks ON and OFF STN DBS and ON and OFF dopaminergic medication. We made three main observations. First, STN DBS increases the anti-saccade error rate while patients are OFF dopamine replacement therapy. Second, there is an interaction between dopamine replacement therapy and STN DBS. More specifically, L-dopa reduces the effect of STN DBS on anti-saccade error rate. Third, STN DBS can induce different effects on pro- and anti-saccades in different patients. These observations provide evidence for an important role for the STN in the circuitry underlying context-dependent modulation of visuomotor action selection.

## Introduction

Neuromodulation or Deep Brain Stimulation (DBS) of several subcortical areas significantly improves motor function in Parkinson’s disease (PD) (Deuschl, Schade-Brittinger et al. 2006, Weaver, Follett et al. 2012). Stimulation of the subthalamic nucleus (STN) improves tremor, drug-induced dyskinesias and motor fluctuations, and may also modulate aspects of executive cognition or emotion (Mallet, Schüpbach et al. 2007, Benabid, Chabardes et al. 2009, Haynes and Haber 2013, Haber and Behrens 2014). However, unlike its motor outcome, the effect of subthalamic DBS on higher order executive functions remains uncertain.

An important component of executive function is inhibitory control: the ability to suppress an impulsive response to external events due to an inappropriate context (Diamond 2013). Dysfunction of inhibitory control is involved in several psychiatric conditions, such as mood disorders, disorders of thought, and addiction (Barkley 1997, Koob and Volkow 2010). In PD, a deficiency of inhibitory control has been reported in different experimental tasks, such as the stop signal task (Gauggel, Rieger et al. 2004), and the Stroop task (Obeso, Wilkinson et al. 2011).

One well-defined and widely used task for evaluating inhibitory control in humans (Guitton, Buchtel et al. 1985, Lueck, Tanyeri et al. 1990, Sereno and Holzman 1995) and non-human primates (Funahashi, Chafee et al. 1993, Gottlieb and Goldberg 1999, Everling and Munoz 2000, Wegener, Johnston et al. 2008) is the oculomotor anti-saccade task. In anti-saccade behavior, unlike pro-saccades, the subject is asked to look away from a target that appears in the peripheral visual field. Inability to suppress the reflexive pro-saccade in the anti-saccade task indicates a deficiency in inhibitory control. Previous work indicates that PD patients generate a larger number of erroneous pro-saccades compared to healthy controls (Briand, Strallow et al. 1999), suggesting that the anti-saccade task may be used to understand how different parts of the basal ganglia network may participate in inhibitory control. The subthalamic nucleus is an important part of the basal ganglia network involved in inhibitory control of a wide variety of sensorimotor, visuomotor and cognitive behaviors (Mink 1996). PD patients with therapeutic implants placed in the subthalamic nucleus may therefore serve as an important model for understanding the role of the STN in the basal ganglia network that modulates anti-saccade performance.

Previous attempts to measure the effect of STN DBS on anti-saccade behavior in PD patients have resulted in variable observations. In two studies PD patients were tested on and off STN DBS for the anti-saccade task while they were on their therapeutic levodopa (L-dopa) dose (Yugeta, Terao et al. 2010, Antoniades, Rebelo et al. 2015). Both studies reported that STN stimulation does not significantly alter the error rate or latency of anti-saccades. In contrast, in a recent study where patients were tested while off L-dopa and equivalents (Goelz, David et al. 2017), STN DBS increased the anti-saccade error rate without changing the latency of correct anti-saccades. The variable results of these three studies calls for a more controlled protocol to examine the effect of STN DBS on anti-saccade behavior. Specifically, since the major difference between the three study designs is the L-dopa state, it is important to evaluate the effect of STN DBS both during the on-and off-medication conditions. We proposed that an interaction between the on-medication state and subthalamic stimulation may potentially explain the conflicting results of previous studies. We measured anti-saccade behavior in a group of PD patients during all four possible conditions of stimulation and medication. Our results show that STN DBS increases anti-saccade error rate. Furthermore, this effect was larger when the patients were OFF L-dopa. In the ON L-dopa condition, the effect of STN DBS on anti-saccade error rate is less pronounced. Therefore, our data supports a complex interaction between stimulation and dopaminergic effects. The subthalamic nucleus appears to be an important part of the circuitry allowing generation of inhibitory anti-saccades in PD. Subthalamic stimulation, which allows for significant improvement in motor function in PD, appears to interfere with frontal inhibitory mechanisms likely generated by the oculomotor loop of the basal ganglia directed to the dorsolateral prefrontal cortex.

## Methods

### Subjects

This study was conducted at the Montreal Neurological Institute, and was reviewed and approved by the ethics committee. 10 PD patients (3 females, mean age 61.2 years old) who had undergone STN DBS surgery were recruited for this study.

### Procedure

Patients were tested in four different combinations of L-dopa or subthalamic stimulation states: 1) OFF L-dopa – ON stimulation, 2) OFF L-dopa – OFF stimulation, 3) ON L-dopa – OFF stimulation, and 4) ON L-dopa – ON stimulation (figure 1a). In each condition, patients were asked to participate in the same series of eye movement tests. Arm tremor was also measured using accelerometers as an index of L-Dopa or STN DBS related effects. Part III of the United Parkinson’s Disease Rating Scale (UPDRS) was performed at the end of the second phase (OFF L-dopa – OFF stimulation) and the fourth phase (ON L-dopa – ON stimulation). (<u>Tremor measurement</u> and <u>Oculomotor tasks</u>). Patients were given a 45-60 minute break between testing in different conditions. The breaks relieved fatigue, and also assured a stable state as patients transitioned between therapeutic conditions. DBS parameters and morning medication doses used in the experiment were identical to those established as therapeutic over time by the patients’ clinicians.

**[FIGURE 1].**
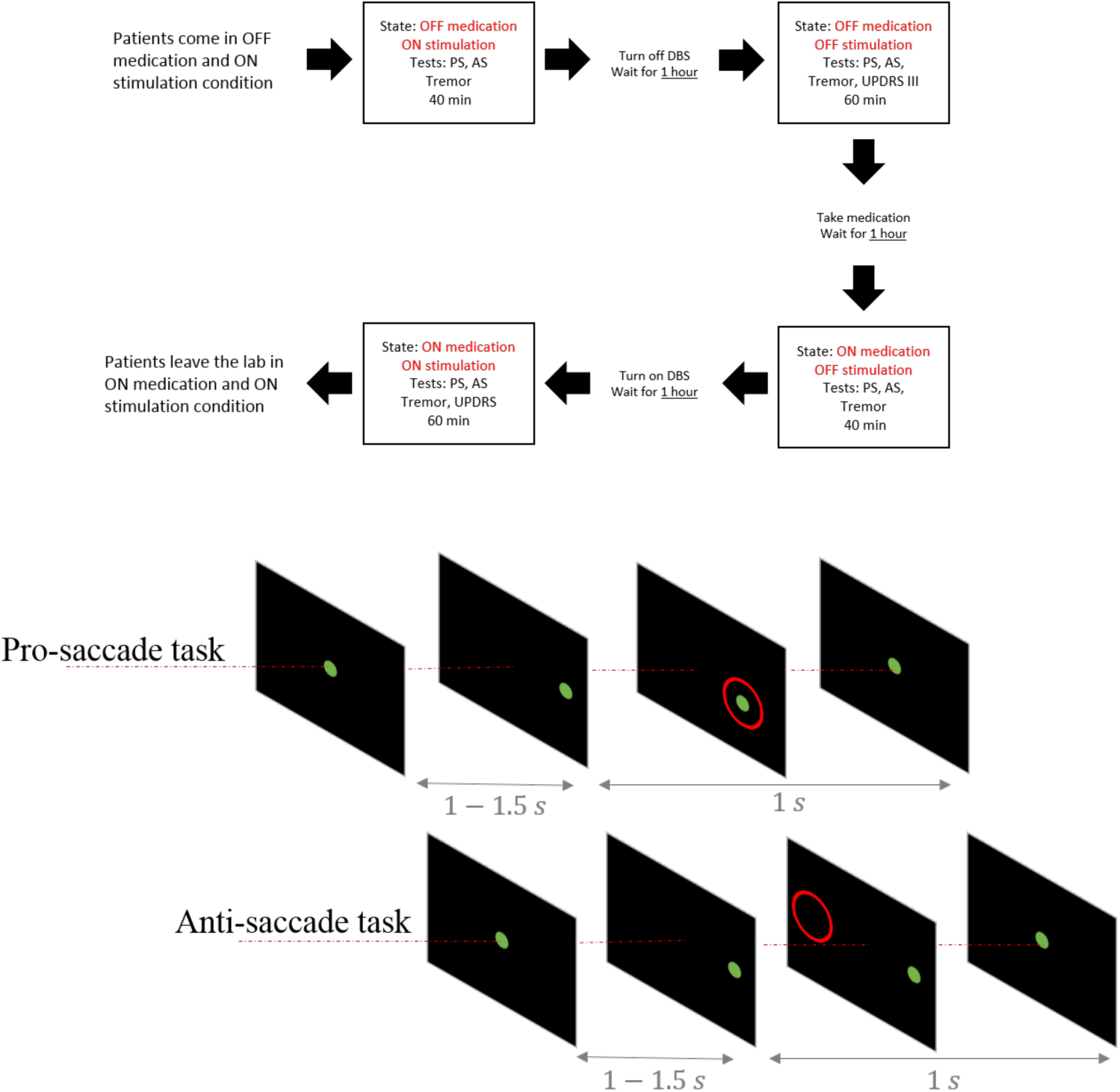
Experiment paradigm and the eye movement task. Top: Each box shows one of the test conditions. Bottom: The schematic figure shows the pro-saccade (top row) and the anti-saccade (bottom row) tasks.

**[FIGURE 2].**
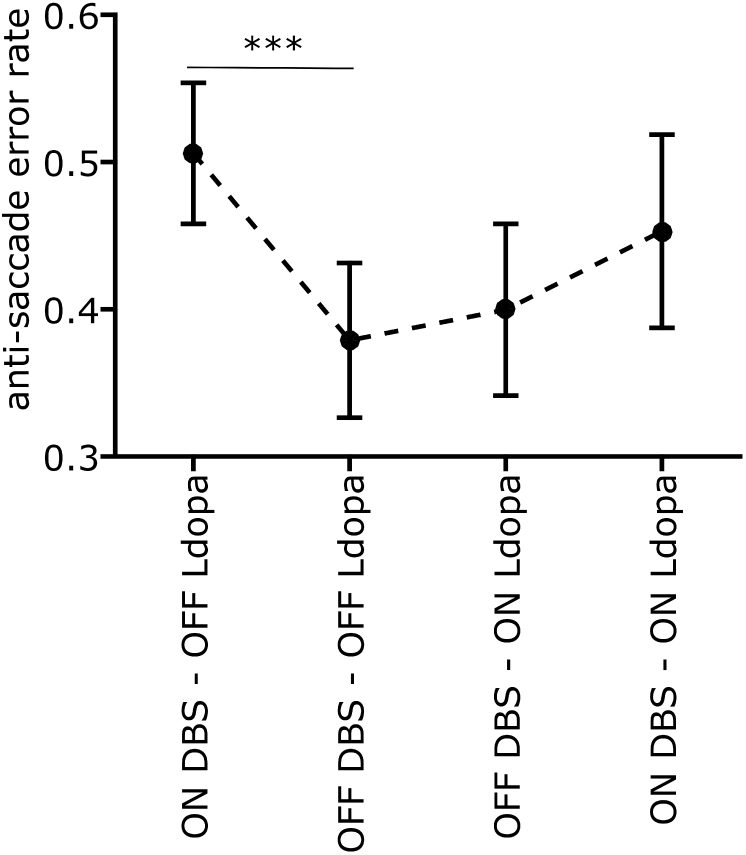
Anti-saccade error rate for different stimulation and L-dopa states. DBS increased the error rate during the off L-dopa condition (*p* < 0.001). Based on our analysis of variance (ANOVA), DBS also increases the error rate while on L-dopa, but not significantly (*p* = 0.15). The L-dopa does not have any significant effect on the error rate on or off DBS (off DBS *p* = 0.32, on DBS *p* = 0.24). The error bars are mean standard errors.

Participants came to the lab early in the morning with their DBS ON, after stopping L-Dopa for about 12 hours. After being given a general introduction to different steps of the protocol that covered the whole day, and full instruction on the eye movement tasks, they went through the first block of tests in the OFF L-dopa – ON stimulation condition. After the first condition, the DBS was turned off, and patients took a 45-60 minute break. Following the break, the same tests were repeated for the OFF L-dopa – OFF stimulation condition. At the end of this condition, patients took their regular dose of L-dopa (table 1) and after the break period, patients were again tested for the same series of tasks in their ON L-dopa – OFF stimulation condition. Eventually, the DBS was turned back on, and after the break, patients were tested in the ON L-dopa – ON stimulation condition.

**[TABLE 1].** Patients’ demographics

| ID | Age | Sex | Years<br>from<br>diagnosis | LED (mg) | UPDRS III<br>DBS OFF | UPDRS III<br>DBS ON |
| --- | --- | --- | --- | --- | --- | --- |
| 1 | 55 | F | 10 | 67 | 65 | 37 |
| 2 | 70 | M | 8 | 133 | 45 | 31 |
| 3 | 63 | M | 17 | 100 | 44 | 30 |
| 4 | 57 | F | 19 | 150 | 54 | 10 |
| 5 | 55 | M | 9 | 100 | 13 | 3 |
| 6 | 70 | M | 17 | 200 | 59 | 35 |
| 7 | 61 | M | 9 | 100 | 56 | 35 |
| 8 | 54 | F | 18 | 266 | 23 | 11 |
| 9 | 62 | M | 7 | 200 | 44 | 29 |
| 10 | 65 | M | 12 | 133 | 21 | 16 |

**LED:** L-dopa equivalent dose (Tomlinson, Stowe et al. 2010). It shows the dose that was given to the patients at the time of the test.

**[TABLE 2].** ANOVA results on anti-saccade error rate

| Name | Estimate | SE | t-stat | DF | p-value | lower | upper |
| --- | --- | --- | --- | --- | --- | --- | --- |
| **db**s | 0.127 | 0.029 | 4.2979 | 36 | 0.0001 (***) | 0.067 | 0.1869 |
| **l**-dopa | 0.0208 | 0.029 | 0.71028 | 36 | 0.48211 | -0.038592 | 0.080193 |
| **db**s * **l**-dopa | -0.0737 | 0.037 | -1.9678 | 36 | 0.05683 | -0.14964 | 0.002257 |

### Tremor measurement

Participants were asked to wear a wrist accelerometer (GENEActiv, Activinsights Ltd., Kimbolton, UK) on their both wrists. They were then seated on a chair with a custom-made arm support. Their hands were supported with the apparatus but free from obstruction. The tremor was measured in this position, with eyes open, for 15 seconds followed by 30 seconds of rest when patients were encouraged to move their hands. This process was repeated three times for each hand.

The accelerometer data was first trend corrected and then preprocessed with Principle Component Analysis (PCA). The first principle component was then used for further analysis. Multi-tapered power spectrum was calculated from the preprocessed data, and the power amplitude for the range of 3-6 Hz was normalized and used as a measure of tremor amplitude.

### Oculomotor tasks

Patients were seated in front of an OLED TV (LG 55EA9800, 122.7 cm × 79.86 cm) at a viewing distance of 57 cm. Chin and forehead rests were used to stabilize their heads and to maximize the accuracy of eye tracking.

Eye position was recorded using a desk-mounted EyeLink 1000 eye tracker (SR Research) at 1000 Hz. Recordings were monocular (right eye only), and the data were analyzed offline.

The eye movement task included two blocks of pro-saccades (60 trials per block) and three blocks of anti-saccades (40 trials per block). The task started with the blocks of pro-saccades and finished with the anti-saccades. Patients had a one minute break between the blocks, and two minutes of break between pro-saccade and anti-saccade trials. For both tasks, the fixation target, a green circle subtending 1.5 visual degrees (luminance 55.2 *cd/m*^2^), appeared on the center of the screen at the beginning of each trial. After 0.5 to 1.5 seconds, it jumped to a 10 degree visual angle on the right or left side of the fixation point. The target then stayed on the screen for 1 second and jumped back to the central position which signaled the beginning of next trial.

In the pro-saccade task, patients were asked to look at the peripheral target as quickly and as accurately as possible. In the anti-saccade task, with the same stimulus, the patients were asked to avoid looking at the peripheral target, and instead to saccade to the opposite side as quickly as possible (figure 1b).

### Data analysis and statistics

Saccades were detected from the recorded eye positions using an automated acceleration-based algorithm (Groh, Born et al. 1997), followed by visual inspection to correct false detections.

For each trial, we extracted parameters related to the corresponding saccade accuracy and speed. In the pro-saccade task, these included saccade latency and amplitude, while for the anti-saccade task, we quantified the latency of correct anti-saccades and error pro-saccades. We also measured the anti-saccade error rate which quantifies the percentage of trials with erroneous pro-saccades during the anti-saccade task. The anti-saccade error rate was used as a measure of inhibitory control failure.

For the pairwise comparisons (*i.e.* comparing one parameter of interest between two conditions), Generalized Linear Mixed Effect models (GLMEs) (Breslow and Clayton 1993), were used to measure the statistical significance of a given difference between the two conditions of interest with the changed factor between the two conditions as the fixed effect (DBS or L-dopa). Due to inter-individual differences in DBS and L-dopa effects, the participants’ identities were introduced as the random effect into the model. For example, in a comparison between the saccade latency in the OFF L-dopa – ON stimulation and OFF L-dopa – OFF stimulation conditions, the stimulation is introduced as the fixed effect, and the patients’ identities as the random effect.

We also used GLME model to run a mixed-effect ANOVA across conditions for different eye movement parameters. For each measurement (e.g. reciprocal latencies and error rates), we used the ‘fitglme’ function in MATLAB to fit the GLME model to the data.

For GLME models to fit the data probability distribution, for all the measurements, the Bayesian Information Criterion (BIC) was used to compare between different models with different output distributions: Normal, Gamma, and Inverse Gaussian. Based on these model comparisons, for the error rate and latencies, we used Normal and Inverse Gamma distributions, respectively.

## Results

To measure the effect of STN DBS on anti-saccade behavior, we tested 10 PD patients (3 females, mean age 61.2 ± 2.97; see table 1) in four different conditions measuring both pro-saccade and anti-saccade behavior. Patients were tested ON or OFF L-dopa, while ON or OFF DBS.

Our tremor measurements indicate that both L-dopa and DBS have significant effect in decreasing tremor amplitude in patients (t-stat = 2.15, p-value = 0.03 for L-dopa, and t-stat = 3.04, p-value = 0.004 for DBS), confirming that effective STN DBS and L-dopa were delivered during the oculomotor task. Moreover, the UPDRS-III evaluations show that the combination of DBS and L-dopa improved the participants’ motor performance (t-stat = 1.97, p-value = 0.057). To make the protocol shorter and prevent further fatigue in patients, we ran the UPDRS evaluations only in two phases of the test: OFF L-dopa – OFF stimulation and ON L-dopa – ON stimulation. Therefore, we are not able to report the effect of individual treatments and their interactions based on the UPDRS.

For the anti-saccade task, we compared four parameters across the four conditions: anti-saccade error rate (ER), correct anti-saccade latency 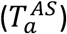, error pro-saccade latency in the anti-saccade task 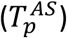, and pro-saccade latency in the pro-saccade task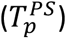.

### Anti-saccade Error Rate

The proportion of erroneous pro-saccades over all the trials is defined as the anti-saccade error rate. Figure (2) shows the mean error rate averaged across patients for four different conditions. During the OFF L-dopa state, turning on STN DBS increases the anti-saccade error rate (t-stat = 5.64, p-value < 0.0001). During the ON L-dopa state, although STN DBS still increases the error rate, this change is smaller and not statistically significant (t-stat = 1.51, p-value = 0.15). There is no effect of L-dopa on the error rate, either ON or OFF DBS.

These observations suggest that STN DBS has a significant effect on anti-saccade error rate, and this effect can be modulated by medication state. To better quantify this interaction, we fitted a mixed-effect generalized linear model (GLME) to the error rate measurements across our cohort of subjects with L-dopa, DBS, and their interaction as the fixed-effects, and the patient identities as the random effect. The results are shown in table (2). Similar to the pairwise comparison above, the GLME analysis indicates that STN DBS increases the error rate, and this effect is statistically significant. Moreover, the interaction analysis indicates that there is a negative interaction between L-dopa and DBS. The results show that the L-dopa decreases the effect of STN DBS on the error rate (t-stat = 1.97, p-value = 0.057). That is, while STN DBS increases the error rate in both L-dopa on and off conditions, this increase is larger while the patients are off L-dopa.

Next, we determined how STN DBS may alter the reaction times during both anti-saccade and pro-saccade tasks.

### Anti-saccade Latency

There are two types of latencies in the anti-saccade task: the latency of correct anti-saccades 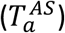, and the latency of the erroneous pro-saccade 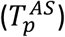. On average, the latency of pro-saccades is shorter than that of the correct anti-saccades, possibly due to further processing needed to generate anti-saccades (Everling and Munoz 2000), generally referred to as the “anti-saccade cost”. Here, we measure the changes in these two types of latencies caused by changes related to being on or off L-dopa, and on or off DBS.

The average latencies for the four conditions are shown in figure (3b). Our results show that the DBS *decreases* the latency for both error pro-saccades (t-stat = 3.12. p-value = 0.006), and correct anti-saccades (t-stat = 2.82, p-value = 0.011). However, L-dopa only *increases* the pro-saccade latency (t-stat = 2.34, p-value = 0.03), and has no significant effect on anti-saccade latency (t-stat = 1.55, p-value = 0.14). Our interaction analysis also shows that there is no interaction between the DBS and L-dopa effects either for pro-saccade (t-stat = 0.62, p-value = 0.54) or anti-saccade latencies (t-stat = 1.3, p-value = 0.2).

In summary, STN DBS *decreases* the correct pro-saccade and erroneous pro-saccade latencies, while L-dopa *increases* only the pro-saccade latency. Figure (4) summarizes the estimated effects of L-dopa and DBS on 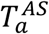 and 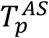. We next consider the effects of interventions on the saccade latency in the pro-saccade task 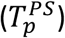.

### Pro-saccade Latency

Figure (3a) shows the changes in the pro-saccade latency 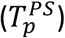 as a function of L-dopa and DBS states. Reproducing previous observations (Temel, Visser-Vandewalle et al. 2008), our results show that STN DBS decreases the pro-saccade latency (t-stat = 3.7, p-value = 0.0007), while L-dopa increases it, though the effect is borderline significant (t-stat = 1.92, p-value = 0.062).

On average, the latency changes that we showed here in the pro-saccade task follow the changes that we reported in previous section for the error pro-saccades in the anti-saccade task (figure 3).

**[FIGURE 3].**
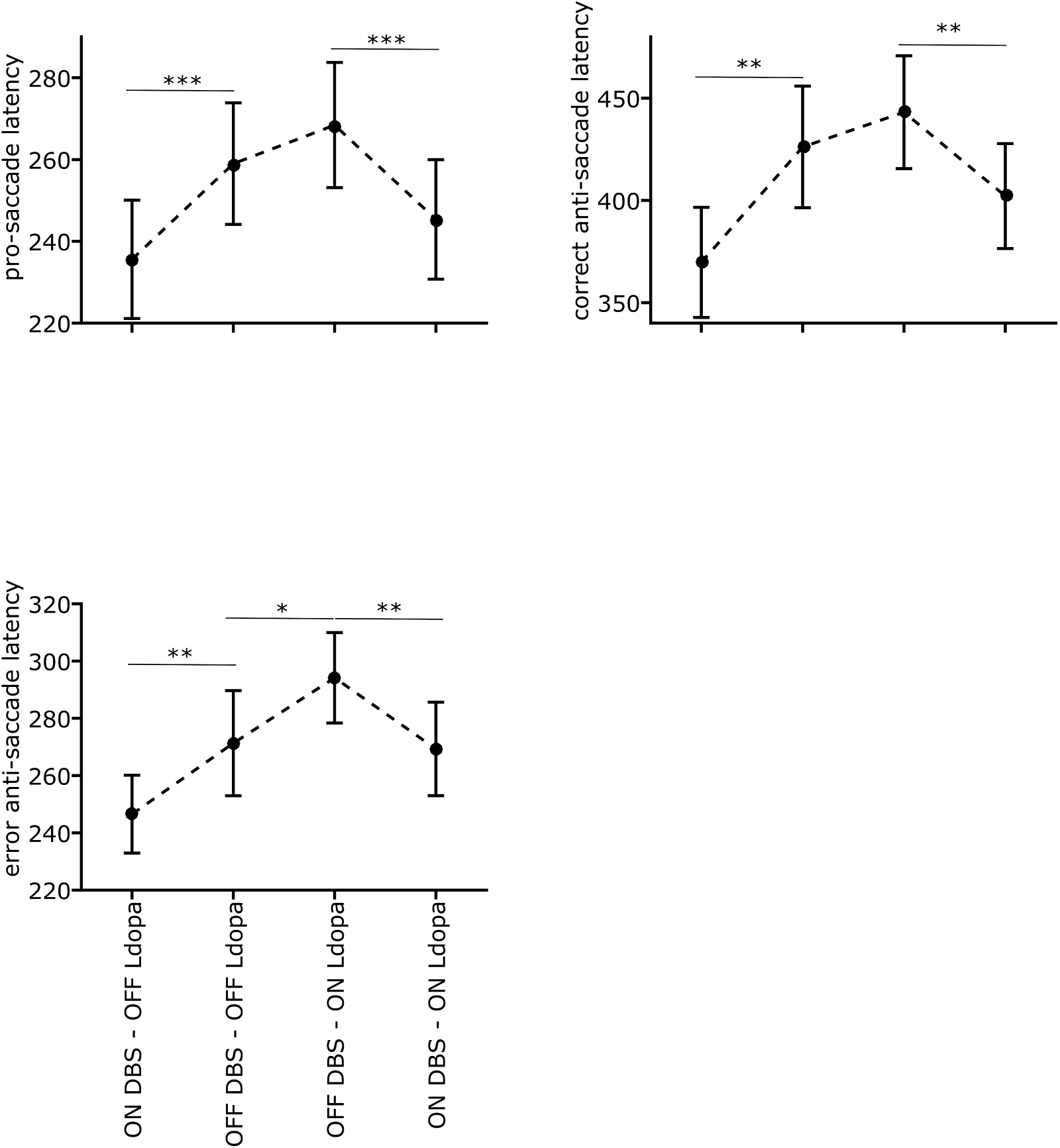
Anti-saccade and pro-saccade latency for different stimulation and L-dopa states. *(Top-left)* The effect of DBS and L-dopa on the pro-saccade latency. Based on our analysis of variance (ANOVA), DBS decreases the pro-saccade latency in both L-dopa conditions (*p* < 0.001). L-dopa increases pro-saccade latency, but it is marginally significance (*p* = 0.062). *(Top-right)* The effect of DBS and L-dopa on the correct anti-saccade latency. DBS decreases the correct anti-saccade latency in both L-dopa conditions (*p* = 0.011). The effect of L-dopa on the correct anti-saccade latency is not statistically significant, but follows the same pattern as in the pro-saccade latency. *(Bottom-left)* The effect of DBS and L-dopa on the erroneous pro-saccade latency in the anti-saccade task. DBS decreases the error pro-saccade latency in both L-dopa conditions (*p* = 0.006), and L-dopa increases the error pro-saccade latency (*p* = 0.03). The error bars are mean standard error.

**[FIGURE 4].**
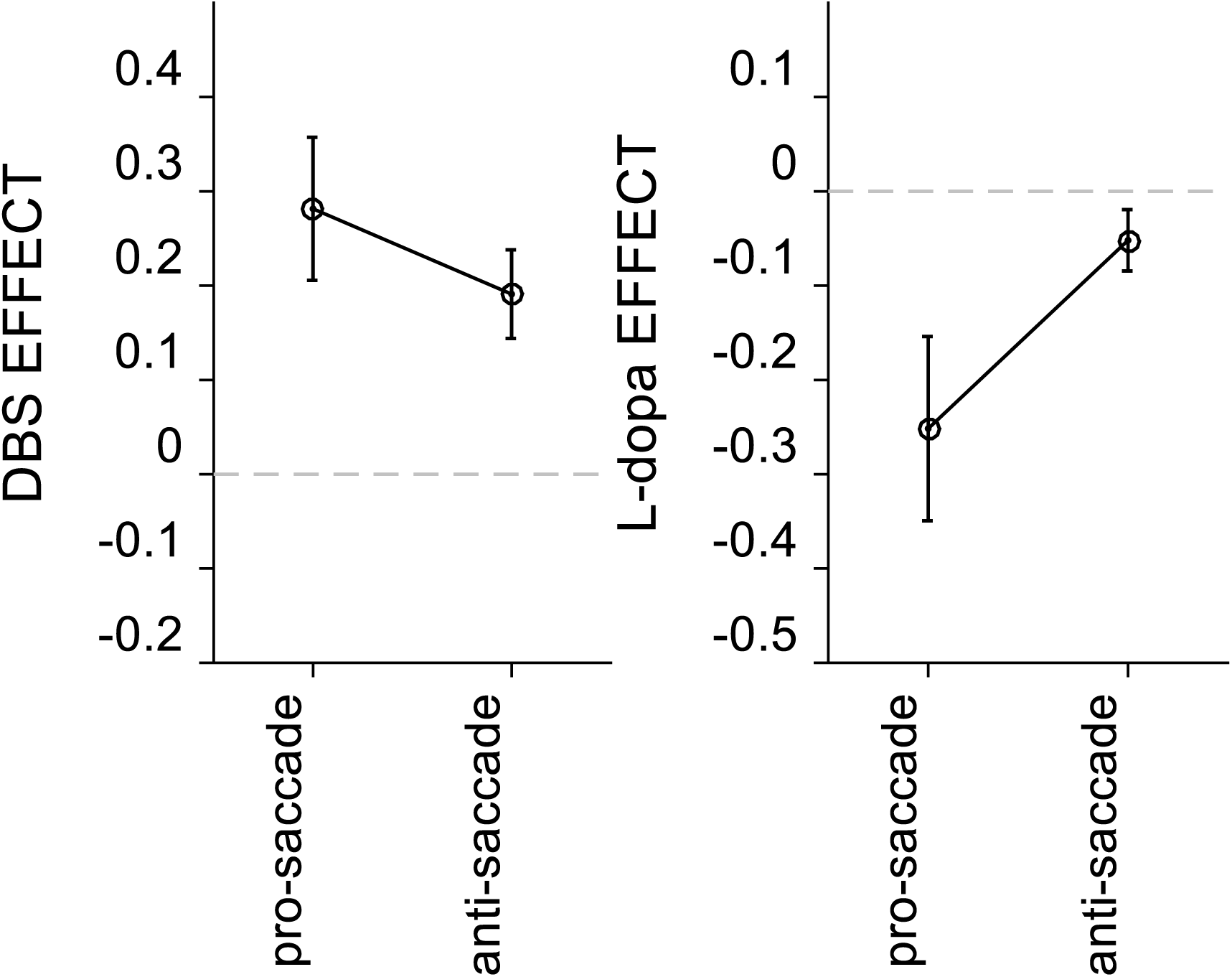
DBS (right) and L-dopa (left) effect on pro-saccade and anti-saccade latencies. Positive and negative effects show decrease and increase in the saccade latencies, respectively. The normalized effect sizes show that DBS decreases both pro-saccade and anti-saccade latencies. The effect is larger in pro-saccade than anti-saccade. L-dopa causes a relatively large increase in pro-saccade latency, but has a close to zero effect on the anti-saccade latency. The error bars are mean standard error.

### DBS Effect on Pro-saccade vs. Anti-saccade Latencies

In figure (5a), we compared the effect of DBS on pro-saccade latencies in the pro-saccade (horizontal axis) and the anti-saccade tasks (vertical axis) within individual subjects. Each circle corresponds to the changes in latencies for each subject. The results show a strong correlation (*r* = 0.6) between these two effects across subjects.

On the other hand, the correlation did not hold when we compared the effect of DBS on pro-saccade and anti-saccade latencies (*r* = 0.07) (figure 5b). The same observation was made with measured pro-saccade latencies in either the pro-saccade task or the anti-saccade. In each subject, therefore, the effect of DBS on the pro-saccade latency is not predictive of its effect on the anti-saccade latency.

**[FIGURE 5].**
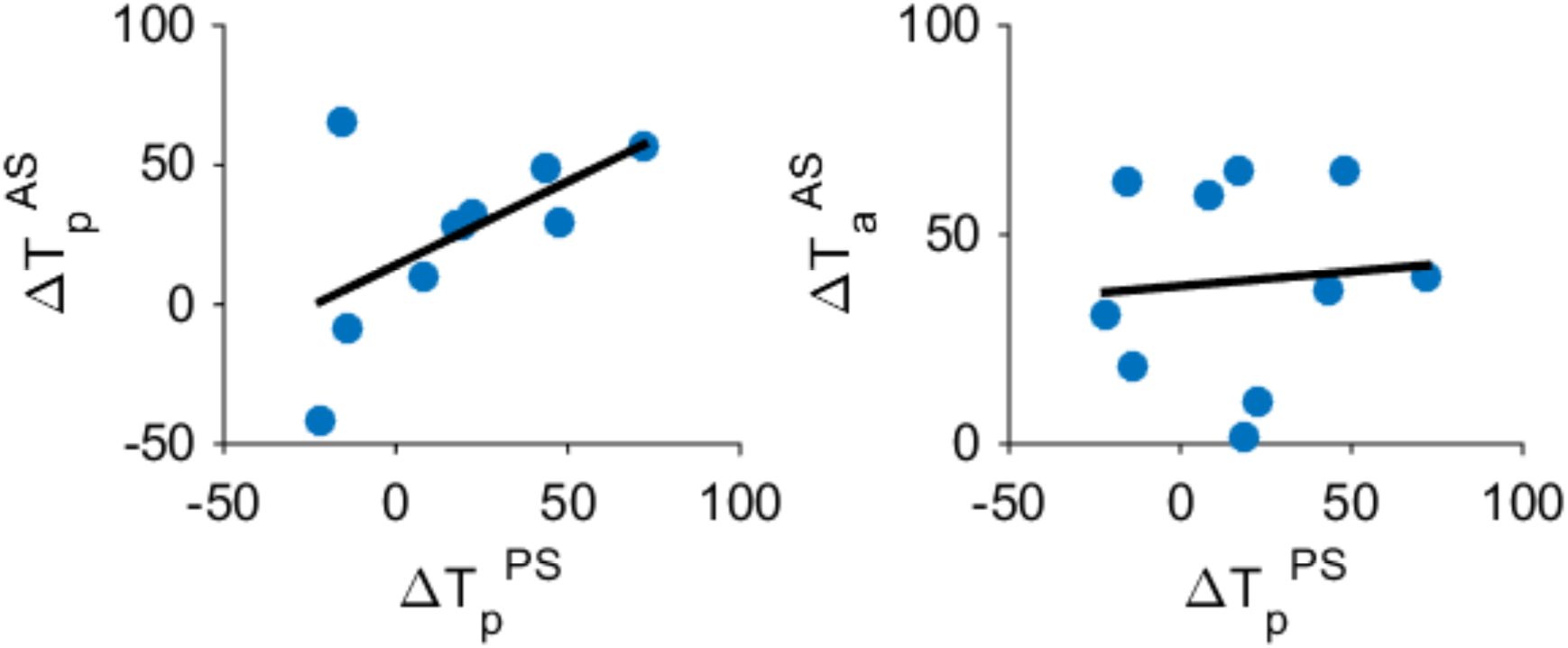
The effect of DBS on pro-vs. anti-saccade latencies. Blue circles show the change in latencies for pro-and anti-saccade in each individual participant. The dark line is the fitter linear model to the data. (Right) There is a strong positive correlation between the change in the pro-saccade latencies in the both pro-saccade and the anti-saccade tasks (*r* = 0.6, *p* < 0.001). (Left) The changes in the pro-saccade latency and the anti-saccade latency caused by DBS are not correlated (*r* = 0.07, *p* = 0.12).

This observation supports the idea that pro-and anti-saccades are controlled by neural circuits that are at least partially independent (see discussion). As shown in this and previous sections, both DBS and L-dopa can differently affect these independent components, and the effect sizes change across subjects.

## Discussion

In the anti-saccade task, two main actions compete to access oculomotor resources: looking towards the target (pro-saccade), and looking away from the target (anti-saccade). Despite overlapping neural pathways (Dias and Segraves 1999), these two inter-related actions are controlled by partially independent circuits (Munoz and Everling 2004). The Superior Colliculus (SC) along with Frontal Eye Fields (FEF) are the main nodes in the oculomotor circuitry that directly project to saccadic premotor circuit to provide the input for saccade initiation.

Therefore, in both pro-saccade and anti-saccade behaviors, the activity of SC and FEF drive the eye movement. However, the ability to suppress a reflexive pro-saccade and perform an anti-saccade reflects successful inhibitory control that is mainly controlled by the frontal and prefrontal areas (Guitton et al., 1985).

In Parkinson’s disease, deficiency in inhibitory control has been reported while patients perform anti-saccade (Kitagawa, Fukushima et al. 1994, Briand, Strallow et al. 1999) and other similar experimental tasks (Cameron, Watanabe et al. 2010). Compared to healthy controls, Parkinson’s disease patients experience more difficulty suppressing the pre-potent reflexive response (e.g. pro-saccade), to generate the voluntary action (e.g. anti-saccade). Although STN DBS and L-dopa successfully reduce the motor symptoms of the disease, their effects on inhibitory control is uncertain.

Here we used the anti-saccade task to measure the effects of STN DBS and dopaminergic medication on inhibitory control in PD patients. The present work demonstrates that L-dopa prolongs only pro-saccade latency, and not anti-saccades. The increase in pro-saccade latency is in keeping with previous work (Michell, Xu et al. 2006), but the effect of L-dopa and dopamine agonists on anti-saccades (error rate and latency) remains controversial (Duka and Lupp 1997, Hood, Amador et al. 2007). The variability of L-dopa dose across patients and heterogeneity of the disease might underlie this uncertainty. Importantly, our results show that the effect of STN DBS on anti-saccade error rate depends on the L-dopa state of the patients. We elaborate more on our findings in the following sections.

### The Effect of STN DBS on Saccades

The subthalamic nucleus, with anatomical connections with both low-level oculomotor nuclei through nigro-collicular projections (Robledo and Féger 1990), and high-level frontal and prefrontal areas (Robledo and Féger 1990, Haynes and Haber 2013) stands at a crossroads in behaviors related to visuo-motor action selection. The precise role of the subthalamic nucleus in different aspects of oculomotor performance in human, especially in inhibitory control, is uncertain, and may be explored using subthalamic stimulation inserted for relief of motor symptoms of PD.

Our results show that STN DBS decreases both pro-saccade and anti-saccade latencies, and increases anti-saccade error rate. On average, our data indicate a larger effect of STN DBS on pro-saccade than anti-saccade latencies (figure 4 – left). Previous attempts at examining the effect of STN DBS on anti-saccade behavior have produced variable observations. Of several studies that specifically addresses effects of STN DBS of anti-saccades (Yugeta, Terao et al. 2010, Antoniades, Rebelo et al. 2015, Goelz, David et al. 2017), only one evaluated patients in their off-L-dopa condition, reporting results similar to the present study (Goelz, David et al. 2017). Two other studies (Yugeta, Terao et al. 2010, Antoniades, Rebelo et al. 2015) tested patients while they were on their regular dose of L-dopa, and reported no significant effect of STN DBS on anti-saccade performance. In the present work, we suggest that the L-dopa state can explain the discrepancies in the previous studies. Based on our results, the STN DBS increases the anti-saccade error rate significantly only when patients are off L-dopa (figure 4 – right). In the on L-dopa condition, STN DBS still increases the error rate but not significantly, which suggests that the effect of STN DBS is mitigated by the L-dopa on state.

Previous electrophysiology, neuroimaging, and computational modeling studies support the role of the STN in cognitive-associational behavior (Temel, Blokland et al. 2005). In particular, more recent electrophysiology studies in animals and humans support the involvement of STN in inhibitory control (Aron and Poldrack 2006, Jahfari, Waldorp et al. 2011, Turner and Pasquereau 2017, Zavala, Jang et al. 2017). The role of the STN in suppressing prepotent responses in inhibitory control tasks (e.g. stop-signal task, Stroop task, and countermanding saccade), has been shown in neuroimaging (Aron and Poldrack 2006), electrophysiology (Schmidt, Leventhal et al. 2013, Turner and Pasquereau 2017), and lesion (Baunez, Nieoullon et al. 1995) studies. In a stop-signal task, for example, Schmidt et al. (2013) showed that STN neurons in rats exhibit short latency responses to the ‘Stop’ signal, which encodes the necessity of canceling a prepared ‘Go’ response. Similar observations were reported in humans using fMRI (Aron and Poldrack 2006). These results are consistent with our observation that STN DBS interferes with inhibitory control, and increases the anti-saccade error rate in PD patients. Moreover, recent studies highlight the importance of beta oscillations in STN and its context-dependent desynchronization for successful motor and cognitive performance (Kondylis, Randazzo et al. 2016, Zavala, Jang et al. 2017). In PD, the pathological hyperactivity (Brown 2003, Crowell, Ryapolova-Webb et al. 2012) and exaggerated beta synchronization affect the normal function of the STN. As suggested by the rate model of direct/indirect pathways of the basal ganglia (DeLong and Wichmann 2007), the STN has an overall inhibitory effect on action generation. Under healthy physiological states, firing of STN neurons selectively suppresses the activity of frontal areas which in turn prolongs action generation. The right level of STN activity is therefore required for generation of contextually appropriate actions, and suppressing inappropriate ones. In PD, the pathological over-expression of synchronized bursts of STN activity leads to abnormally delayed and contextually inappropriate saccades. This may explain the slowness of saccades as well as higher error rate of anti-saccade in PD (Gauggel, Rieger et al. 2004). However, depending on the disease stage, the basal ganglia circuitry is not completely deficient. For example, pathological activity in STN correlates with the symptom severity (Tinkhauser, Pogosyan et al. 2017). Therefore, the residual functionality may still allow some functional control of oculomotor and other associative behaviors, and more pronounced interference with abnormal STN activity may then result in a more detrimental effect on cognitive control. Our results support this hypothesis. Increased inhibition of STN function with DBS may impair a residual control mechanism that allows anti-saccade behavior. Since the STN has an inhibitory effect on saccade-related brain regions (e.g. superior colliculus and frontal eye field), eliminating it from the oculomotor circuit facilitates saccade generation regardless of their importance in the task, which can explain the decreased latencies of both pro-and anti-saccades. Moreover, pro-saccade, since they generally involve a faster process compared to an anti-saccade, would be expected to reach the decision threshold sooner. This may explain the higher error rate that we observe in the ON DBS condition compared to the OFF DBS.

### STN in Oculomotor Circuitry

During a saccade task, previous electrophysiology studies have shown that neuronal firing rates in frontal eye field and superior colliculus ramp up prior to saccade generation (Munoz and Wurtz 1995, Everling and Munoz 2000). The firing rates that correspond to different possible actions in these areas (e.g. pro-saccade vs. anti-saccade) increase to a hypothetical threshold, and the action that first reaches this threshold is selected. The slope of this increase in firing rate determines how fast one commits an eye movement. Therefore, a control mechanism on this slope can favor one action to another based on the context.

Particularly, since the pro-saccade is the incorrect behavior in the anti-saccade task, its corresponding neuronal activity in frontal areas should be controlled to increase less rapidly compared to the anti-saccade. As part of the basal ganglia indirect pathway, STN’s activity can decelerate the increase of firing rates for an action (e.g. pro-saccade in our example) when necessary given the contextual information. Based on our results, we suggest that STN has a crucial role in this coordination process.

Our correlation analysis shows that the DBS effect on pro-saccade in individual subjects does not predict the effect on their anti-saccade (figure 5). These observations support the idea that anti-saccade and pro-saccade behaviors are partially controlled by independent circuits, with frontal and prefrontal areas being more involved in anti-saccade behavior (Munoz and Everling 2004), and the pro-saccades mainly controlled by the superior colliculus. In general, our results indicate that DBS and L-dopa can differentially affect pro-saccade and anti-saccade behaviors. This is in contrast with the assumption in previously proposed models (Noorani and Carpenter 2013).

STN can influence oculomotor behavior through both the nigro-collicular and pallido-thalamic pathways (Robledo and Féger 1990). While both pro-and anti-saccade behaviors are influenced by the two pathways (Everling, Dorris et al. 1999, Everling and Munoz 2000), the pathway that targets the frontal areas (i.e. pallido-thalamic) has more influence on the anti-saccade task, as the frontal areas have shown to be crucially important in correct anti-saccade behavior (Guitton, Buchtel et al. 1985, Funahashi, Chafee et al. 1993, Munoz and Everling 2004). This dual pathway model can explain the previous observation on the differential effect of pallidal and subthalamic DBS on anti-saccade behavior (Antoniades, Rebelo et al. 2015). In their results, Antoniades et al. showed that for patients on L-dopa medication, STN DBS did not have any effect on the anti-saccade error rate. However, stimulation of internal Globus Pallidus (GPi) did decrease the error rate. Unlike STN, GPi has connections with only the frontal areas (and not superior colliculus) through the thalamocortical pathway. Therefore, inhibiting GPi mainly influences the frontal areas which leads to more prominent effect on anti-saccade behavior. On the other hand, STN DBS can have significant effects on superior colliculus as well as frontal areas. In our data, the uncorrelated effects of STN DBS on pro-saccades and anti-saccades (Figure 5) support the dual pathway model of STN DBS effect on oculomotor behavior.

## Summary

In summary, our results support the crucial role of STN in the anti-saccade task. We draw three important conclusions based on our results: first, the pro-saccade and anti-saccade are partially driven by independent mechanisms, and DBS and L-dopa can modulate the two systems independently. Second, eliminating STN from the saccade circuitry decreases the reaction times for both pro-and anti-saccades. As pro-saccade is in general generated faster than anti-saccade, the DBS effect is relatively larger on pro-saccades than anti-saccades, and the action facilitation caused by STN DBS leads to higher anti-saccade error rate. Third, even though STN activity is abnormal in PD, the residual functionality of STN is needed for complex cognitive behaviors, and total elimination of STN by DBS shuts down the control mechanism, and leads to higher error rate.

